# IKKβ signaling mediates metabolic changes in the hypothalamus of a Huntington’s disease mouse model

**DOI:** 10.1101/2021.04.08.438894

**Authors:** Rana Soylu-Kucharz, Ali Khoshnan, Åsa Petersén

**Author notes:** Address for correspondence: Rana Soylu-Kucharz, Translational Neuroendocrine Research Unit, BMC D11, 221 84 Lund, Sweden.

## Abstract

**Background:** Huntington’s disease (HD) is a neurodegenerative disorder caused by a CAG repeat expansion in the huntingtin (*HTT*) gene. Metabolic changes are associated with HD progression, and underlying mechanisms are not fully known. As the IKKβ/NF-κB pathway is an essential regulator of metabolism, we investigated the involvement of IKKβ, the upstream activator of NF-κB in hypothalamus-specific HD metabolic changes.

**Methods:** Using viral vectors, we expressed amyloidogenic N-terminal fragments of mutant HTT (mHTT) fragments in the hypothalamus of mice without IKKβ in the CNS (IKKβ^-/-^) and control mice (IKKβ^+/+^). We assessed effects on body weight, metabolic hormones, and hypothalamic neuropathology.

**Results:** Hypothalamic expression of mHTT led to an obese phenotype only in female mice. CNS-specific inactivation of IKKβ prohibited weight gain in females, which was independent of neuroprotection and microglial activation.

**Conclusions:** The expression of mHTT in the hypothalamus causes metabolic imbalance in a sex-specific fashion, and central inhibition of the IKKβ pathway attenuates the obese phenotype.

## Introduction

Huntington’s disease (HD) is a fatal neurodegenerative disorder caused by a CAG repeat expansion in the huntingtin (HTT) gene (HDCRG, 1993). Although the clinical diagnosis is based on typical motor symptoms, affected individuals also suffer from non-motor symptoms such as cognitive decline, psychiatric symptoms, and metabolic disturbances, which often precede motor symptoms by several years (Bates et al., 2015; Cheong, Gabery, & Petersen, 2019). Metabolic changes in HD include weight loss despite adequate or even higher caloric intake (Aziz et al., 2008; Marder et al., 2009). A higher baseline body mass index (BMI) has been associated with a slower disease progression (van der Burg et al., 2017). Hence, identifying the underlying mechanisms of metabolic changes in HD may reveal novel targets for therapeutic interventions to modify disease progression.

The hypothalamus is a master regulator of metabolism (Cakir & Nillni, 2019; Timper & Bruning, 2017). Imaging studies have identified hypothalamic changes in both prodromal and symptomatic HD patients (Douaud et al., 2006; Kassubek, Gaus, & Landwehrmeyer, 2004; Politis et al., 2008; Soneson et al., 2010). Analyses of postmortem hypothalamic tissue from HD cases and animal models (Cheong et al., 2019) showed loss of neuronal populations expressing orexin (hypocretin), oxytocin, and vasopressin, as well as altered metabolic pathways in several nuclei (Baldo et al., 2019; Gabery, Halliday, Kirik, Englund, & Petersen, 2015; Gabery et al., 2010; Petersen et al., 2005). Inactivation of mutant HTT (mHTT) selectively in the hypothalamus in the transgenic BACHD mouse model prevented developing a metabolic phenotype with obesity accompanied by leptin and insulin resistance (Gray et al., 2008; Hult et al., 2011). Similarly, local expression of mHTT in the hypothalamus using recombinant adeno-associated viral (rAAV) vectors in mice led to hyperphagic obesity with leptin and insulin resistance (Hult et al., 2011; Soylu-Kucharz, Adlesic, Baldo, Kirik, & Petersen, 2015). Transgene expression was present in several appetite-regulating hypothalamic cell populations (i.e., AGRP, POMC)(Hult et al., 2011). As a result, even though these experiments have proved a causal link between hypothalamic expression of mHTT and the development of metabolic imbalance in mice, the mechanisms of hypothalamic mHTT induced metabolic imbalance are still unknown.

The inhibitor of κB kinase β/ nuclear factor-κB (IKKβ/NF-κB) signaling pathway plays a significant role in obesity and overeating and is enriched in the hypothalamus (Meng & Cai, 2011; X. Zhang et al., 2008). Hyperphagia has been shown to activate IKKβ/NF-κB in the hypothalamus through increased endoplasmic reticulum stress, and suppression of IKKβ/NF-κB results in reduced food intake and normalized metabolic phenotype in mice (Douglass, Dorfman, Fasnacht, Shaffer, & Thaler, 2017; X. Zhang et al., 2008; Y. Zhang, Reichel, Han, Zuniga-Hertz, & Cai, 2017). The IKKβ/NF-kB pathway is also activated by mHTT and has been associated with HD pathogenesis (Atwal et al., 2011; Becanovic et al., 2015; Khoshnan et al., 2004; Sarkar et al., 2011; Thompson et al., 2009; Trager et al., 2014). In the R6/1 mouse model of HD, brain-specific deletion of IKKβ impaired the behavioral phenotype and led to exacerbated neurodegeneration with an activated microglial response in the striatum (Ochaba et al., 2019). However, it is not known whether the IKKβ/NF-κB pathway is involved in the development of HD metabolic imbalance. In the present study, our aim was to determine whether inactivation of the IKKβ/NF- κB pathway would prevent hypothalamic-induced metabolic changes induced by mHTT. We therefore performed injections of rAAV vectors expressing mHTT into the hypothalamus of mice without IKKβ in the CNS (IKKβ^-/-^) and compared metabolic effects to control mice with the floxed allele of IKKβ (IKKβ^+/+^).

## Materials and methods

### Animals

The experimental procedures performed on mice were carried out using the approved guidelines in the ethical permit approved by the Lund University Animal Welfare and Ethics committee in the Lund-Malmö region (ethical permit numbers M20-11 and M65-13). Generation of IKKβ^lox/lox^ (referred to as IKKβ^+/+^) mice (Li, Omori, Labuda, Karin, & Rickert, 2003) and Nestin-Cre mice (Betz, Vosshenrich, Rajewsky, & Muller, 1996) was described previously. Nestin/IKKβ^lox/lox^ (referred to as IKKβ^-/-^) mice were generated following several generations of backcrossing. Mice were obtained through crossing IKKβ^-/-^ mice with IKKβ^+/+^ mice. The experiments were carried out on 2-6 months old mice old, both male and female mice, with the genotypes of IKKβ^-/-^ mice and their IKKβ^+/+^ WT littermates. Genotyping for IKKβ^+/+^ was performed using the following primer sequence 5’-GTC ATT TCC ACA GCC CTG TGA-3’ and 5’-CCT TGT CCT ATA GAA GCA CAA C-3’, as described previously (Chen et al., 2003). The animals were kept at 12 hours night/day cycle with free access to a standard chow diet and water.

### Adeno-associated viral vectors

To investigate the effect of the IKK pathway on the development of the metabolic phenotype, we performed stereotactic injections of recombinant adeno-associated viral (rAAV) vectors into the hypothalamus of IKKβ^+/+^ and IKKβ^-/-^ mice. The viral vector was a pseudotyped rAAV2/5 vector (transgene was flanked by two inverted terminal repeats of the AAV2 and packaged in an AAV5 capsid), where the human mHTT gene of 853 amino acids length (853HTT79Q) (Hult et al., 2011). The human Synapsin-1 promoter drove the mHTT gene expression.

### Viral vector injections

The animals were anesthetized by air mask inhalation of isoflurane (2% isoflurane in O2/N2O (3:7)). The mouse head was fixed with a nose clamp and ear bars in the stereotaxic apparatus. Following the head position’s fine-tuning on the stereotaxic frame, the skull was thinned with a dental drill to make a borehole at the determined anterior-posterior and medial-lateral hypothalamic coordinates. Subsequently, the final dorsal-ventral coordinates were measured from the dura mater. The stereotaxic coordinates chosen for hypothalamic injections were: 0.6 mm posterior to bregma, -0.6 mm lateral to the bregma, and 5.2 mm ventral to the dura mater, selected according to the mouse brain atlas (Franklin & Paxinos, 2008).

The rAAV vector delivery was performed bilaterally in the hypothalamus. The silica glass capillary (with an outer diameter of ∼ 80 μm) attached to a 5 μl Hamilton syringe (Nevada, USA) was used for the virus delivery. A total volume of 0.5 μl viral vector solution was pulled in a glass capillary, and the capillary was descended from dura mater to target coordinates slowly. At the target, the first 0.1 μl of the total volume was injected. After 30 seconds, the viral suspension was delivered at a rate of 0.05μl/15s until the whole volume was delivered. To allow the brain to absorb the viral vector solution, the capillary was left at the target for an additional 5 minutes at the end of the injection. The viral vector concentration used in the study was 2,1E+14 genome copies (GC)/ml.

### Perfusion and serum collection

To induce deep anesthesia in mice, a terminal dose of pentobarbital (600 mg/kg, Apoteksbolaget) was injected intraperitoneally. The thoracic cavity was opened to expose the heart, and blood was collected from the right ventricle with the 16G needle. Subsequently, a small incision was made to insert a 12-gauge perfusion needle at the tip of the left ventricle. First, the vessels were rinsed with the saline solution at a rate of 10-12 ml/min for a minute, and then it was switched to freshly prepared 4% paraformaldehyde (PFA) ∼0°C for 8 minutes. Following that, the animals were decapitated, and the brains were isolated. The brains were placed in 4% PFA for 24 hours at 4°C for post-fixation. Next, the PFA was replaced with 25% sucrose solution at 4°C for cryoprotection (∼24 hours).

Finally, the fixed brains were sectioned coronally on a semi-automated freezing microtome (Microm HM 450) at 30 μm thick slices and in six series. Until further processing, the brain sections were stored in antifreeze solution (30% glycerol and 30% ethylene glycol in phosphate buffer) at −20 °C.

### Metabolic Measurements

Serum insulin and leptin concentrations were assessed in blood collected from the heart left ventricle prior to perfusion. The blood was kept at room temperature for 30 minutes to clot and spun for 15 minutes at 2500 g. The serum (supernatant) was stored at -80 C. Serum levels of insulin (Crystal Chem Inc, Catalog #90080) and leptin (Crystal Chem Inc, Catalog #90030) were assessed using ELISA according to the manufacturer’s instructions.

### Immunohistochemistry

The free-floating brain sections were used for immunohistochemistry (IHC). All the washing steps (10 minutes/wash) and incubations were performed using gentle agitation on a shaker at room temperature. The sections were washed three times in Tris-buffered-saline (TBS) in 1% Triton X (TBS-T) to remove the antifreeze solution. The endogenous peroxidase activity was blocked by 30 minutes of incubation in 10% methanol with 3% H2O2 in TBS. Following that, the sections were washed three times for 10 minutes in TBS-T. The sections were incubated with 5% serum and bovine serum albumin (BSA) in TBS-T for 1 hour to reduce nonspecific binding of the primary and secondary antibodies. Next, the sections were left in the respective primary antibody solutions in 3% appropriate corresponding serum in TBS-T (anti-huntingtin (sc-8767; 1:500; goat; Santa Cruz), anti-ubiquitin (1:2000; rabbit; Dako), anti-orexin (1:4000; rabbit; Phoenix Pharmaceuticals), anti-tyrosine hydroxylase (1:2000; rabbit; Pel-Freez), anti-GnRH (1:3000, anti-rabbit, Abcam #ab5617), anti-iba-1) and left on shaker for overnight incubation at room temperature. Next, sections were washedthree3 times for 10 minutes in TBS-T. The secondary antibody incubation was performed in 3% respective serum or BSA with TBS-T for 1 hour at room temperature, and the sections were washed three times for 10 minutes in TBS before 3,3′-diaminobenzidine (DAB) development (Vectastain, ABC kit). Brain sections were mounted on chromatin-gelatin coated glass slides. The air-dried sections were left in distilled water for 1 minute and dehydrated in increasing ethanol solutions (70%, 95%, 99%). Finally, the samples were cleared in xylene and covered with glass coverslips using DPX mounting medium (Sigma-Aldrich).

### Stereological analyses

To estimate the numbers of cells positive for orexin, TH, GnRH (in the anterior hypothalamus; AHA), and size of the inclusions, we applied unbiased stereological quantification principles by using the optical dissector method (West, Slomianka, & Gundersen, 1991). Stereological analyses were performed with a Nikon 80i microscope, which is equipped with an X–Y motorized stage (Märzhauser, Wetzlar) and a high precision linear encoder (Heidenhain, Traunreut). The position of the stage and the input from the digital camera were controlled by a computer. The sampling interval was adjusted to count at least 100 cells for each hypothalamus to minimize the coefficient of error. The region of interest was delineated under the 4X objective, whereas the counting was performed using a 60X NA 1.4 Plan-Apo oil objective with a random start systematic sampling routine (NewCast Module in VIS software; Visiopharm A/S, Horsholm). The border delineation processes for orexin, TH, GnRH cell populations were defined by the natural contours of cell populations. The number and size of iba-1 positive microglia/macrophages were quantified in the anatomical borders of the mediobasal hypothalamus. The number of small, medium and large size HTT inclusions was quantified with HTT staining (sc-6787). The inclusions ∼ 0.04 - 0.1 μm size considered small, 0.15 - 0.25 μm medium and ∼ 0.25-0.5 μm large size inclusions. The same size of the area was quantified for all genotypes and gender under blinded conditions. The mean number of assessed inclusions per brain was 755, median 663 with a standard deviation of error 528 inclusions.

### Metabolic tests

For all animals used in the study, body weight was measured bimonthly. Serum insulin and leptin concentrations were assessed in serum. Blood was collected from the heart left ventricle at 18 weeks post-injection, and they were kept at room temperature for 30 minutes to clot, spun for 15 minutes at 2500 g. Serum (supernatant) was aliquoted and stored at -80 C. Serum levels of leptin (Crystal Chem Inc, Catalog #90030) and testosterone (Demeditec, Cat.-No.: DEV9911) were determined with ELISA according to the manufacturer’s instructions.

### Statistical analysis

All statistical analyses were performed using Prism 8 software (GraphPad). The data was initially tested with D’Agostino & Pearson omnibus normality test for normal distribution. Following that, the data was either subjected to Kruskal–Wallis followed by Dunn’s multiple comparison tests or an unpaired t-test with equal SD. The statistical test results and the type of analysis used for each experiment are specified in detail in the results section and figure legends. Statistically significant differences were considered for p<0.05.

## Results

### Inhibition of the IKKβ pathway protects from hypothalamic mHTT-induced obesity in female mice

To assess the effects of mHTT expression on the development of a metabolic phenotype, we injected IKKβ^+/+^ (homozygous for the floxed allele of IKKβ, control group) and IKKβ^-/-^ (expressing Cre-recombinase under nestin promotor) mice with rAAV vectors expressing mHTT (AAV5-853HTT79Q vectors are referred to as HD elsewhere) in the hypothalamus of both sexes. Consistent with our previous findings (Baldo, Soylu, & Petersen, 2013; Hult et al., 2011; Soylu-Kucharz et al., 2015), expression of mHTT in female control mice led to an obese phenotype (**Figure 1A, C)**. The body weight at 18 weeks post-injection was significantly higher in the IKKβ^+/+^HD mice (n=16 and mean= 55.6g, SD=10.2) compared to IKKβ^-/-^HD mice (n=20 and mean=40.6g, SD=9.3, p<0.0001) and uninjected mice of the two genotypes (uninjected IKKβ^+/+^: n=15 and mean=43.7g, SD=7.4, p=0.0026; uninjected IKKβ^-/-^: n=13 and mean=37.1g, SD=9.8, p<0.0001). Hence, inactivation of the IKKβ pathway in Nestin-expressing cells protected female mice from hypothalamic mHTT induced obesity **(Figure 1A)**. The brain-specific deletion of IKKβ did not affect the circulating insulin and leptin levels in mice (Meng & Cai, 2011). Therefore, in this study, we assessed the serum levels of insulin and leptin levels only in mice expressing mHTT in the hypothalamus. In line with the prevention of body weight gain, IKKβ^-/-^HD mice displayed significantly lower serum levels of insulin and leptin than IKKβ^+/+^HD mice **(Figure 1D)**.

**Figure 1:**
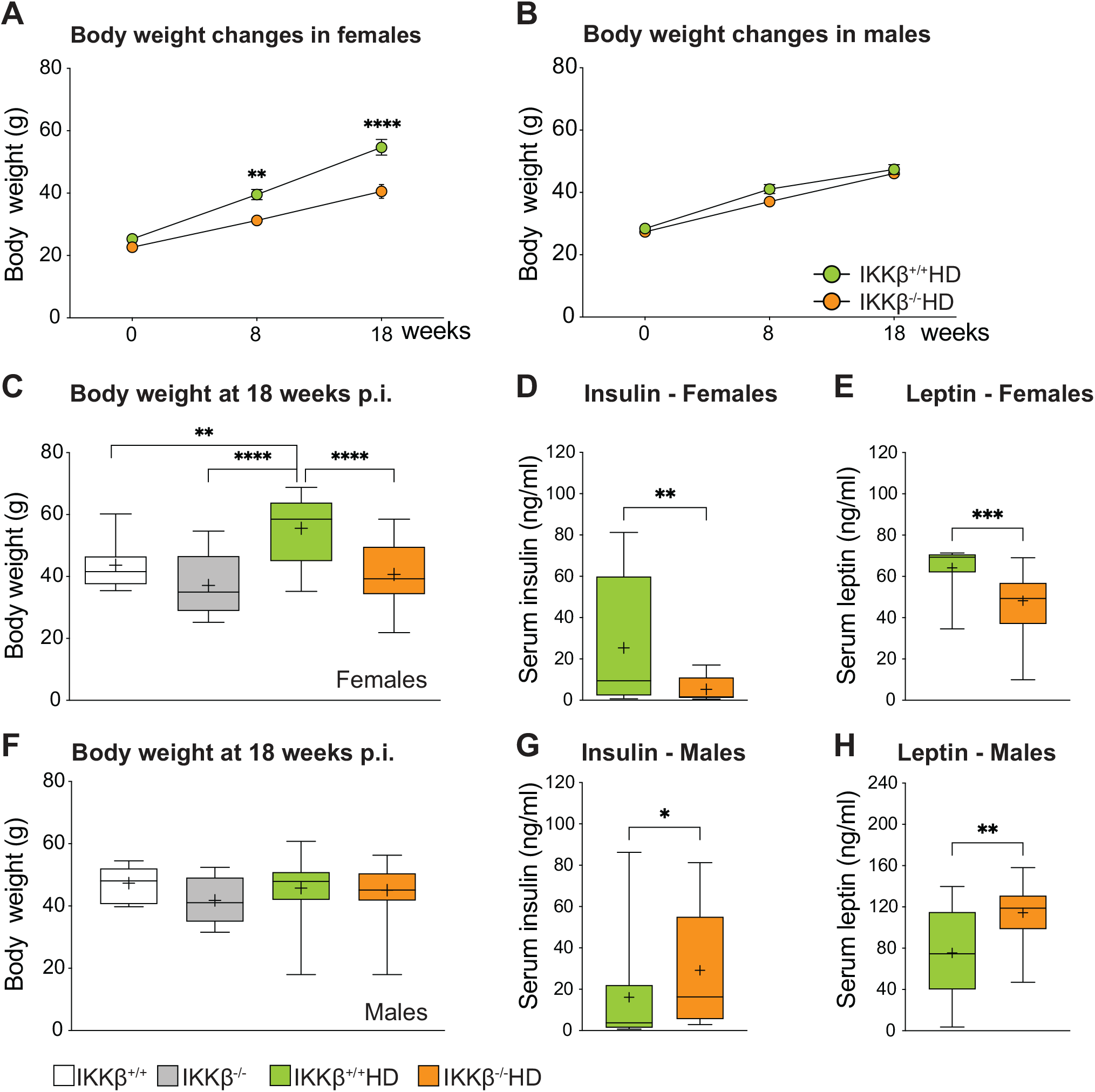
Inactivation of the IKKβ pathway inhibits the development of obesity-induced by mHTT expression in the hypothalamus of female mice. IKKβ^+/+^ and IKKβ^-/-^ mice were injected bilaterally into the hypothalamus with AAV-HTT853-79Q vectors and assessed using metabolic analyses. **(A)** Female IKKβ^+/+^ mice develop increased body weight after hypothalamic injections of AAV5-HTT853-79Q vectors which is prevented in IKKβ^-/-^ female mice (two-way repeated measures ANOVA, effect of time F (2, 62) = 8.920, p<0.0001; effect of genotype F (1, 31) = 20.83, p<0.0001; effect of genotype x time F (2, 62) = 8.920, p=0.0004; followed by a Sidak’s multiple comparisons test: p= 0.0026 at 8 weeks and p< 0.0001 at 18 weeks). **(B)** Male IKKβ^+/+^ and IKKβ^-/-^ mice do not develop obesity after AAV5-HTT853-79Q vector injections (two-way repeated measures ANOVA, effect of time F (2, 72) = 190.4, P < 0.0001; effect of genotype F (1, 36) = 2.359, P = 0.1333; effect of genotype x time F (2, 72) = 1.446, P = 0.2422, followed by a Sidak’s multiple comparisons test). **(C)** Body weight changes at 18 weeks post-injection in females (One-way ANOVA, effect of treatment F (3, 60) = 11.69, P<0.0001 followed by Sidak’s multiple comparisons test). Serum **(D)** insulin (two-tailed, unpaired t-test, n=16/19, p=0.0082) and **(E)** leptin (two-tailed, Mann Whitney test, n=16/19, p=0.0003) concentrations measured by ELISA in females at 18 weeks post injection. **(F)** Body weight changes at 18 weeks post-injection in males (One-way ANOVA, effect of treatment F (3, 55) = 0.7312”, P=0.5378 followed by Sidak’s multiple comparisons test). Serum **(G)** insulin (two-tailed, Mann Whitney test, n=15/21, p=0.0448) and **(H)** leptin (two-tailed, unpaired t-test, n=15/21, p=0.0023) assessments at 18 weeks. Data are represented as box and whisker plots (25– 75 percentile (boxes), min to max (whiskers), median (horizontal line), mean (+)).

IKKβ did not affect the body weight in male mice with or without injections of AAV5-853HTT79Q as assessed up to 18 weeks post-injection **(Figure 1B, F)**. However, analyses of serum levels of insulin and leptin showed that even though there was no effect on body weight, levels of insulin and leptin were significantly elevated in IKKβ^-/-^HD mice compared to IKKβ^+/+^HD mice **(Figure 1G and 1H)**.

### IKKβ is not involved in the mHTT-mediated loss of orexin and TH positive cell populations in HD mice

The development of the metabolic phenotype in HD has been associated with expression of mHTT in the hypothalamus (Hult et al., 2011; Soylu-Kucharz et al., 2015). Here we tested whether the rescue of the metabolic phenotype observed in IKKβ^-/-^HD mice was due to the preservation of metabolism-regulating neuronal populations known to be affected in HD (Cheong et al., 2019). In female mice, there was no benefit of IKKβ silencing as the number of orexin positive cells was comparable in IKKβ^+/+^HD (cell loss ∼64%) and IKKβ^-/-^HD groups (cell loss ∼82%), and both groups had a significantly lower number of orexin cells compared to uninjected groups of female mice **(Figure 2A and 2B)**. Male IKKβ^-/-^HD mice had significantly lower orexin and A13 TH positive cells than IKKβ^+/+^HD mice in the hypothalamus **(Supplementary Figure 1A and 1B)**.

**Figure 2:**
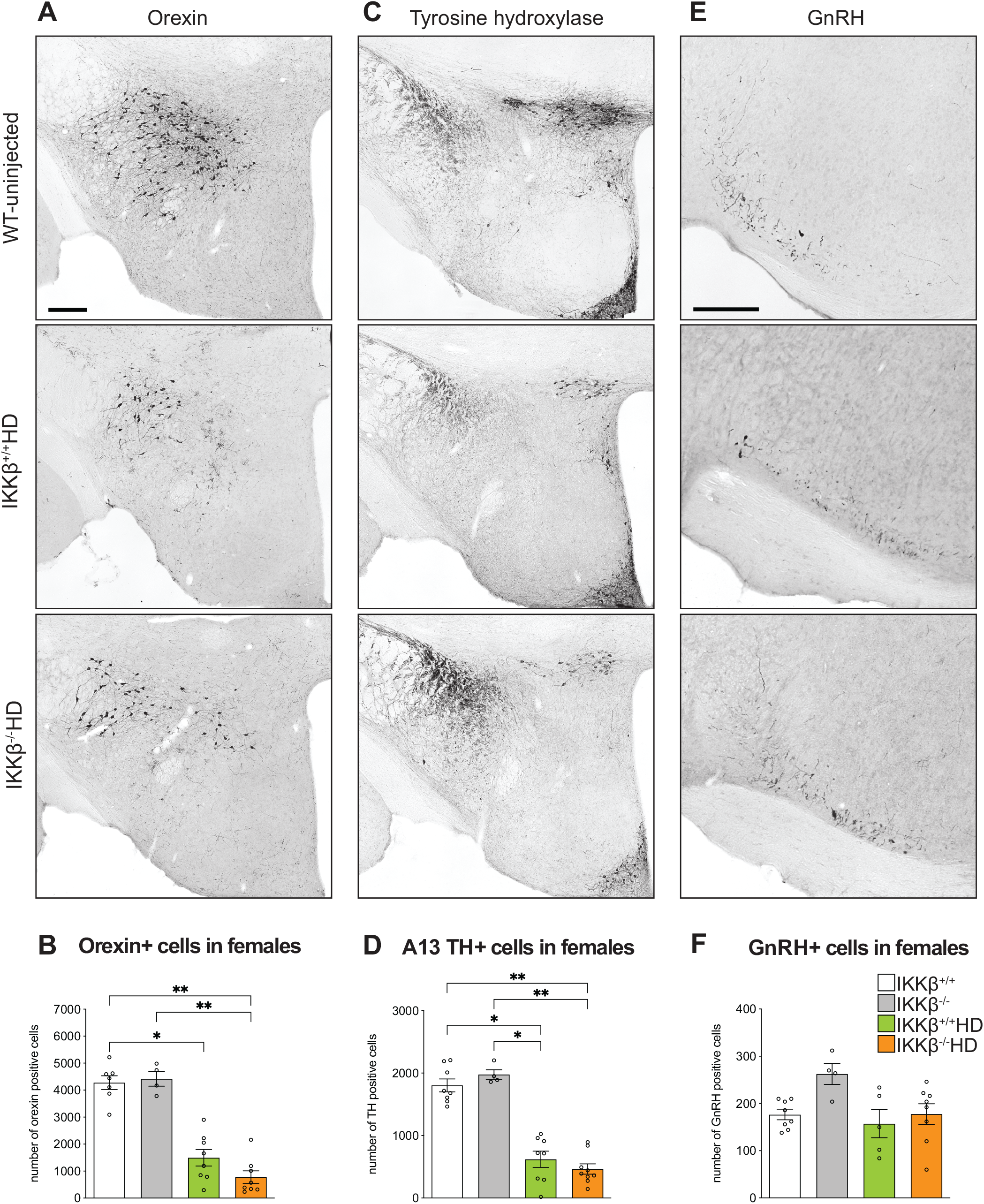
Quantitative analysis of the neuronal populations expressing orexin, TH in the A13 area, and GnRH in female mice at 18-week post-injection of AAV-HTT853-79Q vectors. **(A) Representative** immunohistochemically stained sections show orexin immunopositive cells in the hypothalamus. **(B)** Stereological analysis of orexin immunopositive cells in female mice at the 18 weeks time point (Kruskal-Wallis test followed by Dunn’s multiple comparisons test p=0.0002; n=4-8/group). **(C)** Representative photomicrographs show the A13 TH immunopositive cell population in the hypothalamus. **(D)** Numbers of A13 TH positive cells in the hypothalamus’s zona incerta area (Kruskal-Wallis test followed by Dunn’s multiple comparisons test p=0.0001; n=4- 9/group). **(E)** Representative photomicrographs illustrate the GnRH positive cells in the anterior hypothalamic area of the hypothalamus. **(F)** Stereological quantification of GnRH positive cells in the anterior hypothalamic area (Kruskal-Wallis test followed by Dunn’s multiple comparisons p=0.0802; n=4-8/group). Points on scatter graphs represent total cell count for individual mice, the lines are means, and the whiskers indicate ± SEM. Scale bars represent 200 μm.

The loss of A13 TH positive cells was ∼80% in IKKβ^-/-^HD and ∼65% in IKKβ^+/+^HD when compared to WT and IKKβ^-/-^ uninjected female mice **(Figure 2C and 2D)**.

Reduced testosterone levels are associated with increased circulating levels of leptin and insulin, even in the absence of increased BMI (Luukkaa et al., 1998; Pitteloud et al., 2005). The hypothalamic-pituitary-gonadal (HPG) axis, which regulates testosterone production, is altered in HD (Bird, Chiappa, & Fink, 1976; Kalliolia et al., 2015; Markianos, Panas, Kalfakis, & Vassilopoulos, 2005; Papalexi et al., 2005; Petersen & Bjorkqvist, 2006; Saleh et al., 2009; Soylu-Kucharz, Baldo, & Petersen, 2016; Van Raamsdonk et al., 2007). The number of GnRH positive cells was comparable **(Figure 2F and Supplementary Figure 1C)**; however, the total circulating level of testosterone was diminished in IKKβ^-/-^HD male mice by 60% compared to IKKβ^+/+^HD male mice expressing mHTT in the hypothalamus **(Supplementary Figure 1D)**.

### The number or size of Iba-1 positive microglial cells in the mediobasal hypothalamus

Given that IKKβ is one of the mediators of microglial activation and energy balance (Karin, 1999; Liu, Zhang, Joo, & Sun, 2017), we investigated whether protection from hypothalamic mHTT induced obesity in IKKβ^-/-^ female mice was due to alteration in microglial activation. Nonetheless, the number and size of Iba1 positive cells in the mediobasal hypothalamus of both IKKβ^+/+^HD and IKKβ^-/-^HD female mice were comparable **(Figure 3A-C)**.

**Figure 3:**
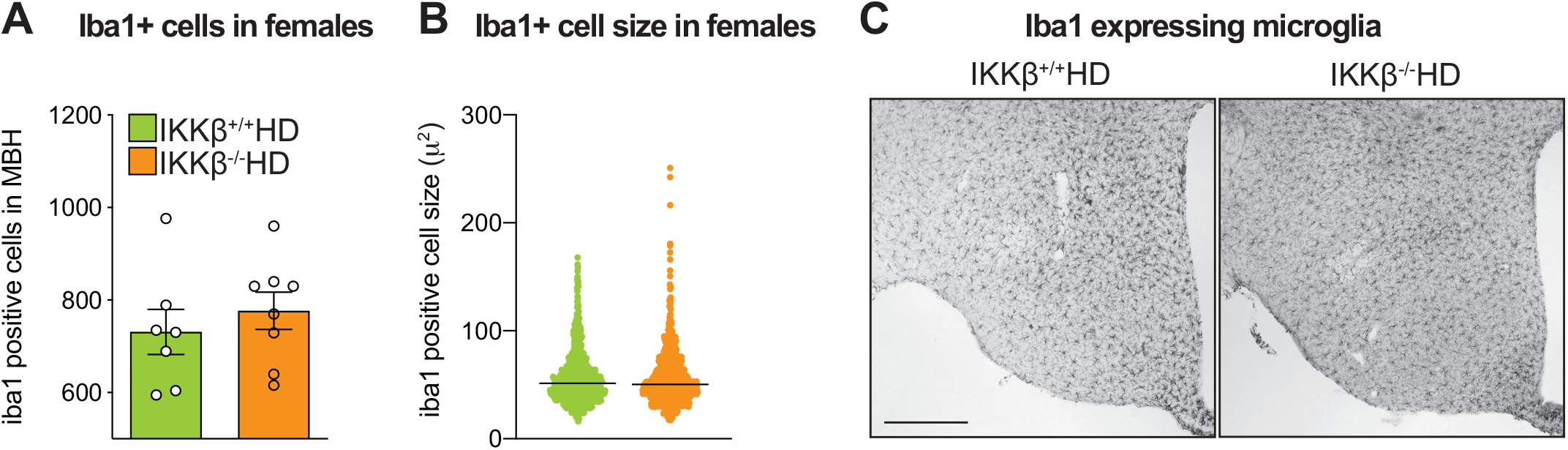
No effect of the IKKβ pathway on the degree of iba-1 positive cell activation at 18 weeks post-injection in females. Stereological assessment of **(A)** the total number of iba-1 positive cells (two-tailed, unpaired t-test, n=7/8, p=0.477) and **(B)** the size of Iba-1 positive cells (two-tailed, Mann-Whitney test, n=7/8 animals, n=783/911 cells/genotype, p=0.9601) in the mediobasal hypothalamus (MBH) 18 weeks after injections of AAV5-HTT853-79Q vectors in IKKβ^+/+^ and IKKβ^-/-^ mice. In (A), data are represented as scatter dot plots, and bars represent mean ± SEM, and in (B), data are represented as scatter dot plots, and lines represent median.

### IKKβ^*-/-*^HD female mice display an increased number of small-sized inclusions of mHTT

Reduction of IKKβ activity decreases the cleavage of both WT and mHTT and prevents the accumulation of mHTT inclusions (Khoshnan, Ko, Tescu, Brundin, & Patterson, 2009; Thompson et al., 2009). IKKβ silencing studies also showed impaired clearance of mHTT and worsening HD pathological phenotypes in vivo and vitro (Khoshnan et al., 2009; Thompson et al., 2009). In our model, inclusions were increased in IKKβ^-/-^HD compared to IKKβ^+/+^HD in both female and male mice **(Figure 4A-C)**. The small size inclusions were responsible for the increase, as the number of medium or large size inclusions were similar between IKKβ^+/+^HD and IKKβ^-/-^HD groups **(Figure 4D-F)**. Altogether, these results show the increase in inclusion formation correlates with previous reports on IKKβ silencing in HD (Criollo et al., 2010; Khoshnan et al., 2009; Ochaba et al., 2019).

**Figure 4:**
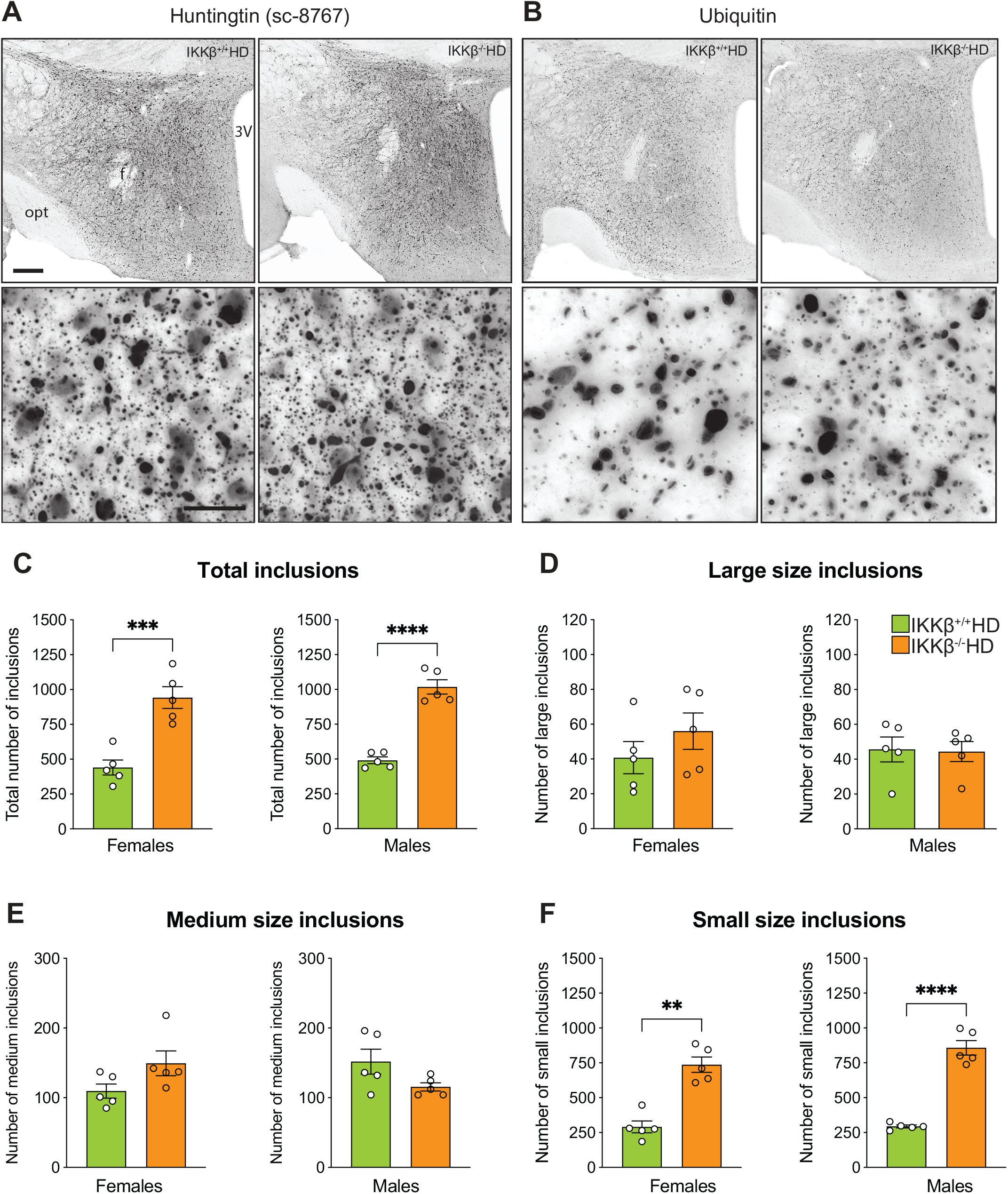
Inactivation of the IKKβ pathway leads to increased numbers of huntingtin inclusions in the hypothalamus. Representative photomicrographs of sections processed for immunohistochemistry for **(A)** huntingtin (using the sc-8767 antibody) and **(B)** ubiquitin demonstrating the formation of inclusions in the hypothalamus after injections of AAV-HTT853-79Q vectors in IKKβ^+/+^ and IKKβ^-/-^ mice. Stereological quantification of sections processed with the huntingtin antibody show **(C)** the total number of inclusions (Females: two-tailed, Mann-Whitney test, n=5/group, p=0.0008; Males: two-tailed, Mann-Whitney test, n=5/group, p<0.0001), (D) large-sized inclusions (Females: two-tailed, Mann-Whitney test, n=5/group, p=0.3095; Males: two-tailed, unpaired t-test, n=5/group, p=0.8992), **(E)** medium-sized inclusions (Females: two-tailed, unpaired t-test, n=5/group, p=0.0635); Males: two-tailed, unpaired t-test, n=5/group, p=0.0911) and **(F)** small-sized inclusions (Females: two-tailed, Mann-Whitney test, n=5/group, p=0.0079; Males: two-tailed, unpaired t-test, n=5/group, p<0.0001) inclusions in female and male mice at 18 weeks post-injection. Points on scatter graphs represent total inclusion count for individual mice, the lines are means, and the whiskers indicate ± SEM. Scale bars represent 200 μm and 25 μm on lower and higher magnifications, respectively.

## Discussion

Alterations in energy metabolism may affect disease progression in HD as weight loss is part of the clinical phenotype, and a higher BMI has been associated with the slower clinical decline (van der Burg et al., 2017). Understanding the underlying biological cause of metabolic disturbances in HD may unravel novel therapeutic targets for this fatal neurodegenerative disorder. The IKKβ/NF- κB pathway has been implicated in HD pathogenesis (Khoshnan & Patterson, 2011), but it has not been investigated in the context of HD metabolic and hypothalamic alterations. Here, we expressed mHTT in the hypothalamus of control mice and compared effects to mice with IKKβ inactivated in the CNS. We found that hypothalamic mHTT expression induces obese phenotype selectively in the female mice and inactivation of the IKKβ prevents it. Gender differences play a role in HD as the severity and rate of the motor symptoms progression has been suggested to be faster in women than men with HD (Zielonka et al., 2013; Zielonka et al., 2018; Zielonka & Stawinska-Witoszynska, 2020). Previous studies indicated that sex also affects the HD metabolic and behavioral manifestation in animal models (Dorner, Miller, Barton, Brock, & Rebec, 2007; Sjogren et al., 2019; Soylu-Kucharz et al., 2016). Even though the expression of mHTT in the hypothalamus of male mice did not affect body weight, male IKKβ^-/-^HD displayed high serum leptin and insulin levels. As testosterone deficiency is associated with metabolic syndrome exemplified by increased circulating leptin levels and insulin resistance, it is possible that the increase in serum leptin and insulin levels could be due to reduced circulating testosterone levels in IKKβ^-/-^HD mice.

Obese phenotype can occur due to increased caloric intake, decreased activity, metabolic rate, or combination of these factors. Previously, we demonstrated that the obese phenotype caused by hypothalamic mHTT expression was due to hyperphagia as general motor activity and basal metabolic rate were unaltered in these mice (Hult et al., 2011). The mice in this study were housed in a separate animal unit that lacks behavior testing platforms and limiting the number of in-house cages. Therefore, we were not able to re-test basic parameters such as food intake and locomotor activity. However, as we have previously shown that obese phenotype was caused by increased food intake, we speculate that the silencing of the IKKβ expression prohibited hyperphagia-induced obesity in HD mice. The orexin and A13 TH neuronal populations are involved in metabolism regulation (Adeghate et al., 2020; Shi et al., 2013) and they are affected in HD (Gabery et al., 2010; Hult et al., 2011; Petersen et al., 2005; Soylu-Kucharz et al., 2015). As the inactivation of the IKKβ signaling did not affect the preservation of these cells, we can speculate that orexin and A13 TH neuropathology are not the central cell populations responsible for the development of HD bodyweight alterations.

In conclusion, our study shows that hypothalamic expression of mHTT leads to a metabolic imbalance in a sex-specific fashion. The weight gain phenotype induced by the mHTT in female mice is prevented by IKKβ inactivation and is independent of orexin TH neuroprotection, and microglial activation.

## Author contributions statements

RS, ÅP, and AK conceived and designed the experiments. RSK performed the experiments. RSK and ÅP analyzed the data. RSK and ÅP wrote the first draft of the manuscript. All authors were involved in editing the manuscript and approved the final version.

## Figure legends

**Supplementary figure 1:**
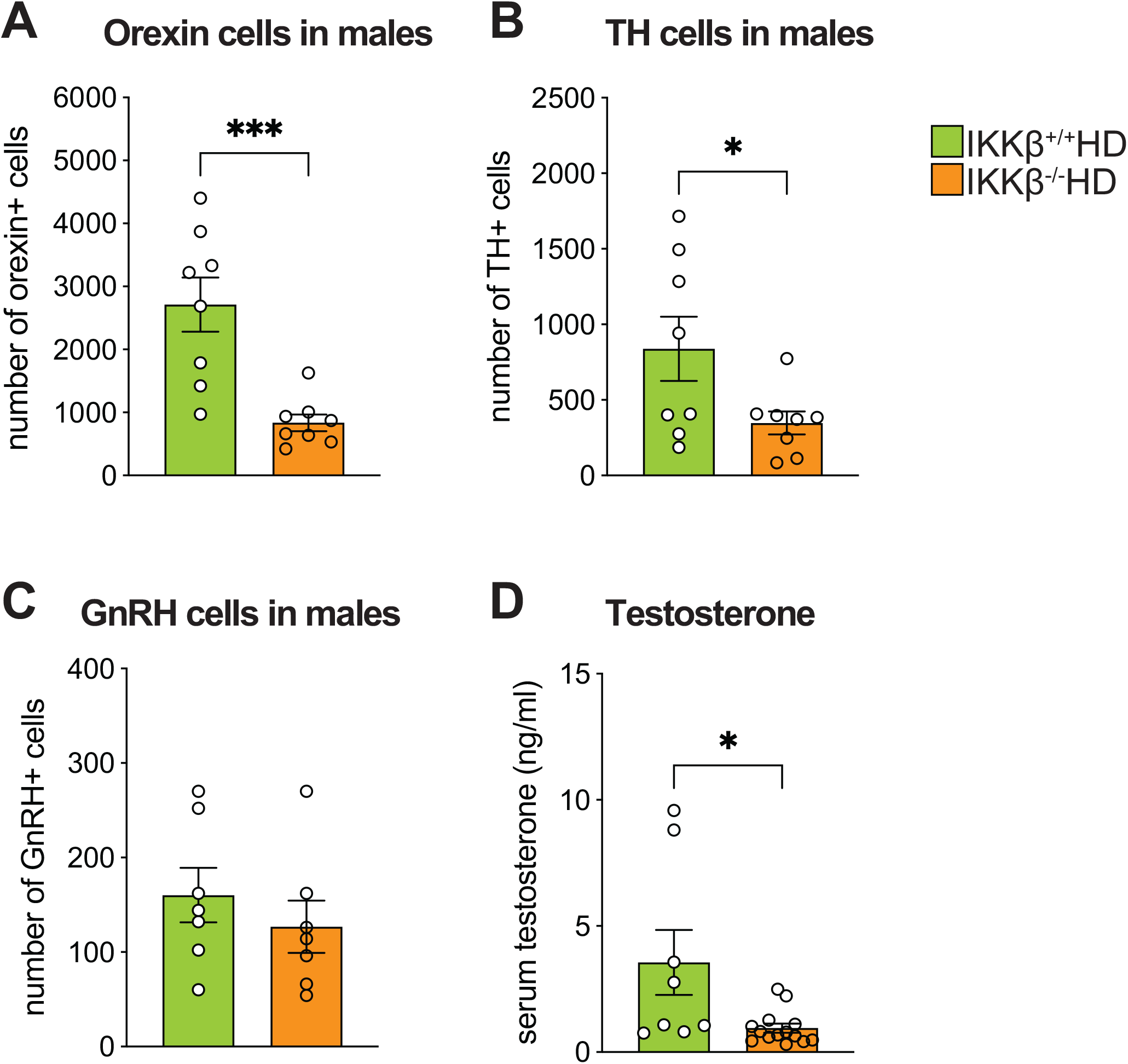
Effects on the orexin, GnRH, and A13 TH cell populations in the hypothalamus in male mice after injections of AAV-HTT853-79Q vectors. Stereological quantification of **(A)** orexin (two-tailed, unpaired t-test, n=8/group, p=0.001), **(B)** TH (two-tailed, unpaired t-test, n=8/group, p=0.0471) and **(C)** GnRH (two-tailed, unpaired t-test, n=7/group, p=0.3648) positive cells in IKKβ^+/+^ and IKKβ^-/-^ males expressing mHTT in the hypothalamus at 18 weeks post-injection. **(D)** Changes in serum testosterone levels in male IKKβ^+/+^ and IKKβ^-/-^ mice at 18 weeks post-injection of AAV-853HTT-79Q in the hypothalamus (two-tailed, Mann-Whitney test, n=8/14, p=0.0159). Data are represented as scatter dot plots, the lines are means, and the whiskers indicate ± SEM.

## Acknowledgments

This work was supported by grants from the Swedish Medical Research Council (grant numbers 2013/03537 and 2018/02559), the Province of Skåne State Grants (ALF) as well as the Knut and Alice Wallenberg Foundation (# 2019.0467) to ÅP. ÅP is a Wallenberg Clinical Scholar (Knut and Alice Wallenberg Foundation # 2019.0467). RSK was supported by Svenska Sällskapet för Medicinsk Forskning fellowship. We are grateful for the excellent technical assistance provided by Björn Anzelius, Anneli Josefsson, Ulla Samuelsson and Ulrika Sparrhult-Bjork at Lund University.

